# The apoptotic, inflammatory, and fibrinolytic actions of vasoinhibin are in a motif different from its antiangiogenic HGR motif

**DOI:** 10.1101/2023.08.18.553934

**Authors:** Juan Pablo Robles, Magdalena Zamora, Jose F. Garcia-Rodrigo, Alma Lorena Perez, Thomas Bertsch, Gonzalo Martinez de la Escalera, Jakob Triebel, Carmen Clapp

## Abstract

Vasoinhibin is a proteolytic fragment of the hormone prolactin that inhibits blood vessel growth (angiogenesis) and permeability, stimulates the apoptosis and inflammation of endothelial cells and promotes fibrinolysis. The antiangiogenic and antivasopermeability properties of vasoinhibin were recently traced to the HGR motif located in residues 46 to 48, allowing the development of potent, orally active, HGR-containing vasoinhibin analogs for therapeutic use against angiogenesis-dependent diseases. However, whether the HGR motif is also responsible for the apoptotic, inflammatory, and fibrinolytic properties of vasoinhibin has not been addressed. Here, we report that HGR-containing analogs are devoid of these properties. Instead, the incubation of human umbilical vein endothelial cells with oligopeptides containing the sequence HNLSSEM, corresponding to residues 30 to 36 of vasoinhibin, induced apoptosis, the nuclear translocation of NF-κB, the expression of genes encoding leukocyte adhesion molecules (*VCAM1* and *ICAM1*) and proinflammatory cytokines (*IL1B, IL6, TNF*), and the adhesion of peripheral blood leukocytes. Also, the intravenous or intra-articular injection of HNLSSEM-containing oligopeptides induced the expression of *Vcam1, Icam1, Il1b, Il6, Tnf* in the lung, liver, kidney, eye, and joints of mice and, like vasoinhibin, these oligopeptides promoted the lysis of plasma fibrin clots by binding to plasminogen activator inhibitor-1 (PAI-1). Moreover, the inhibition of PAI-1, urokinase plasminogen activator receptor, or NF-κB prevented the apoptotic and inflammatory actions. In conclusion, the functional properties of vasoinhibin are segregated into two different structural determinants. Because apoptotic, inflammatory, and fibrinolytic actions may be undesirable for antiangiogenic therapy, HGR-containing vasoinhibin analogs stand as selective and safe agents for targeting pathological angiogenesis.

## Introduction

The formation of new blood vessels (angiogenesis) underlies the growth and repair of tissues and, when exacerbated, contributes to multiple diseases, including cancer, vasoproliferative retinopathies, and rheumatoid arthritis (1). Antiangiogenic therapies based on tyrosine kinase inhibitors (2,3) and monoclonal antibodies against vascular endothelial growth factor (VEGF) or its receptor (4) have proven beneficial for the treatment of cancer and retinal vasoproliferative diseases (5). However, disadvantages such as toxicity (6–8) and resistance (9) have incentivized the development of new treatments.

Vasoinhibin is a proteolytically generated fragment of the hormone prolactin that inhibits endothelial cell proliferation, migration, permeability, and survival (10). It binds to a multi-component complex formed by plasminogen activator inhibitor-1 (PAI-1), urokinase plasminogen activator (uPA), and the uPA receptor on endothelial cell membranes, which can contribute to the inhibition of multiple signaling pathways (Ras-Raf-MAPK, Ras-Tiam1-Rac1-Pak1, PI3K-Akt, and PLCγ-IP_3_-eNOS) activated by several proangiogenic and vasopermeability factors (VEGF, bFGF, bradykinin, and IL-1β) (10). Moreover, vasoinhibin, by itself, activates the NF-κB pathway in endothelial cells to stimulate apoptosis (11) and trigger the expression of inflammatory factors and adhesion molecules, resulting in leukocyte infiltration (12). Finally, vasoinhibin promotes the lysis of a fibrin clot by binding to PAI-1 and inhibiting its antifibrinolytic activity (13).

The antiangiogenic determinant of vasoinhibin was recently traced to a short linear motif of just three amino acids (H46-G47-R48) (HGR motif) which led to the development of heptapeptides comprising residues 45 to 51 of vasoinhibin that inhibited angiogenesis and vasopermeability with the same potency as whole vasoinhibin (14) (Figure 1a). The linear vasoinhibin analog (Vi45-51) was then optimized into a fully potent, proteolysis-resistant, orally active cyclic retro-inverse heptapeptide (CRIVi45-51) (Figure 1a) for the treatment of angiogenesis-dependent diseases (14). Noteworthy, thrombin generates a vasoinhibin of 48 amino acids (Vi1-48) that contains the HGR motif (Figure 1a). Vi1-48 is antiangiogenic and fibrinolytic (15), suggesting that the HGR motif could also be responsible for the apoptotic, inflammatory, and fibrinolytic properties of vasoinhibin. This possibility needed to be analyzed to support the therapeutic future of the HGR-containing vasoinhibin analogs as selective and safe inhibitors of blood vessel growth and permeability. Moreover, the identification of specific functional domains within the vasoinhibin molecule provides insights and tools for understanding its overlapping roles in angiogenesis, inflammation, and coagulation under health and disease.

**Figure 1.**
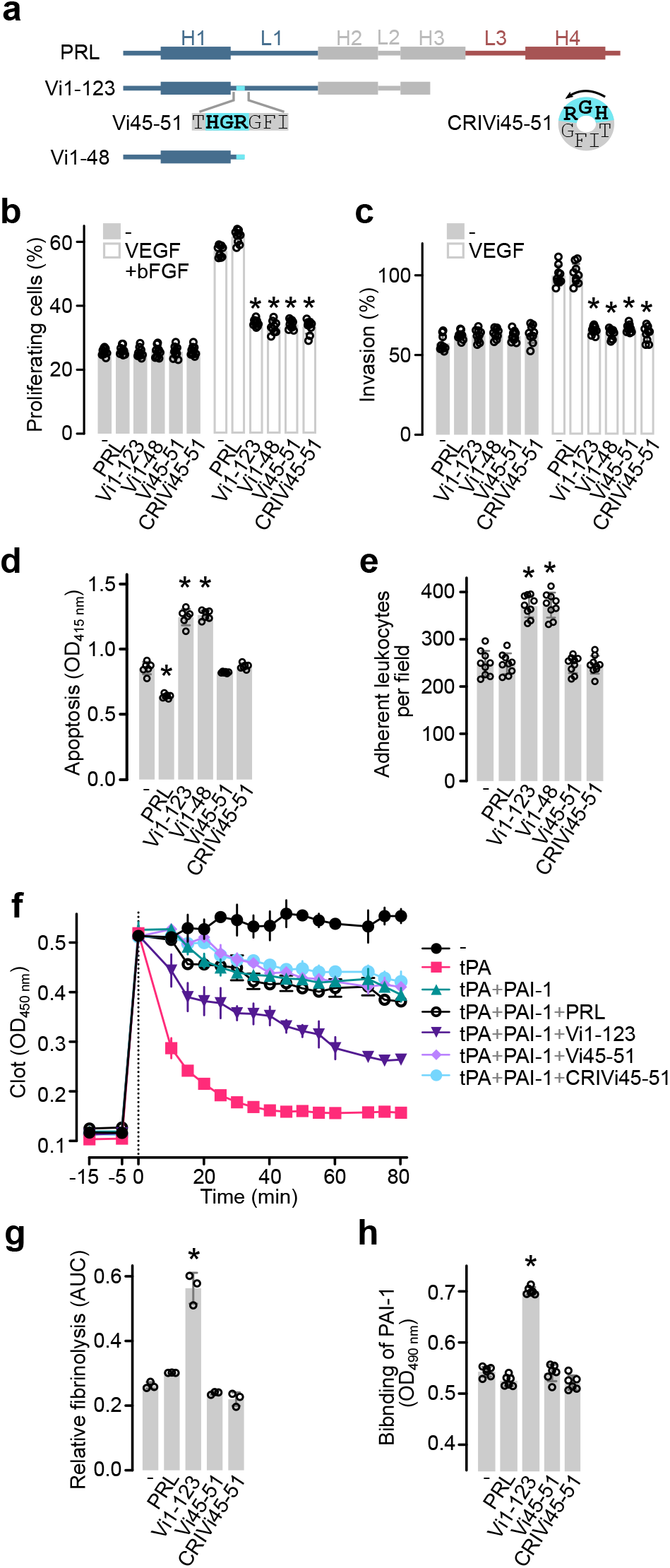
HGR-containing vasoinhibin analogs are neither inflammatory, apoptotic, nor fibrinolytic. **(a)** Diagram of the secondary structure of prolactin with 199 residues (PRL), the 123-residue (Vi1-123), and the 48-residue (Vi1-48) vasoinhibin isoforms. The location and sequence of the linear HGR-containing vasoinhibin analog (Vi45-51) and the cyclic retro-inverse HGR-containing vasoinhibin analog (CRIVi45-51) are illustrated. The antiangiogenic HGR motif is highlighted in cyan. (**b**) Effect of 100 nM PRL, Vi1-123, Vi1-48, Vi45-51, or CRIVi45-51 on the proliferation of HUVEC in the presence or absence of 25 ng mL^−1^ VEGF and 20 ng mL^−1^ bFGF. Values are means ± SD relative to total cells (*n* = 9). **(c)** Effect of 100 nM PRL, Vi1-123, Vi1-48, Vi45-51, or CRIVi45-51 on the invasion of HUVEC in the presence or absence of 25 ng mL^−1^ VEGF. Values are means ± SD relative to VEGF stimulated values (*n* = 9). \**P* <0.0001 vs. VEGF+bFGF (-) or VEGF (-) controls (Two-way ANOVA, Dunnett’s). **(d)** Apoptosis of HUVEC in the absence (-) or presence of 100 nM of PRL, Vi1-123, Vi1-48, Vi45-51, or CRIVi45-51 (*n = 6*). **(e)** Leukocyte adhesion to a HUVEC monolayer in the absence (-) or presence of 100 nM of PRL, Vi1-123, Vi1-48, Vi45-51, or CRIVi45-51 (*n = 9*). **(f)** Lysis of a plasma clot by tissue plasminogen activator (tPA) alone or together with the plasminogen activator inhibitor-1 (PAI-1) in the presence or absence of PRL, Vi1-123, Vi45-51, or CRIVi45-51 (*n = 3*). **(g)** Fibrinolysis relative to tPA and tPA+PAI-1 calculated with the area under the curve (AUC) of **(f)**. **(h)** Binding of PAI-1 to immobilized PRL, Vi1-123, Vi1-48, Vi45-51, or CRIVi45-51 in an ELISA-based assay (*n = 6*). Individual values are shown with open circles. Values are means ± SD, \**P* <0.001 vs. control without treatment (-) (One-way ANOVA, Dunnett’s).

## Materials and Methods

### Reagents

Six linear oligopeptides (>95% pure) acetylated and amidated at the N- and C-termini, respectively (Table 1), the linear (Vi45-51), and the cyclic-retro-inverse-vasoinhibin-(45–51)-peptide (CRIVi45–51) were synthesized by GenScript (Piscataway, NJ). Recombinant vasoinhibin isoforms of 123 (Vi1-123) (16) or 48 residues (Vi1-48) (15) were produced as reported. Recombinant human PRL was provided by Michael E. Hodsdon (17) (Yale University, New Haven, CT). Human recombinant plasminogen activator inhibitor 1 (PAI-1) was from Thermo Fisher Scientific (Waltham, MA) and human tissue plasminogen activator (tPA) from Sigma Aldrich (St. Louis, MO). Rabbit monoclonal anti-PAI-1 [EPR17796] (ab187263, RRID:AB_2943367) and rabbit polyclonal anti-β-tubulin antibodies (Cat# ab6046, RRID:AB_2210370) were purchased from Abcam (Cambridge, UK), and mouse monoclonal anti-uPAR (RRID:AB_2165463) from R&D systems (Minneapolis, MN, Cat# MAB807, RRID:AB_2165463). The NF-κB activation inhibitor BAY 11-7085 and lipopolysaccharides (LPS) from *<u>Escherichia coli</u>* O55:B5 were from Sigma Aldrich. Recombinant human vascular endothelial growth factor-165 (VEGF) was from GenScript, and basic fibroblast growth factor (bFGF) was donated by Scios, Inc. (Mountain View, CA).

**Table 1.**
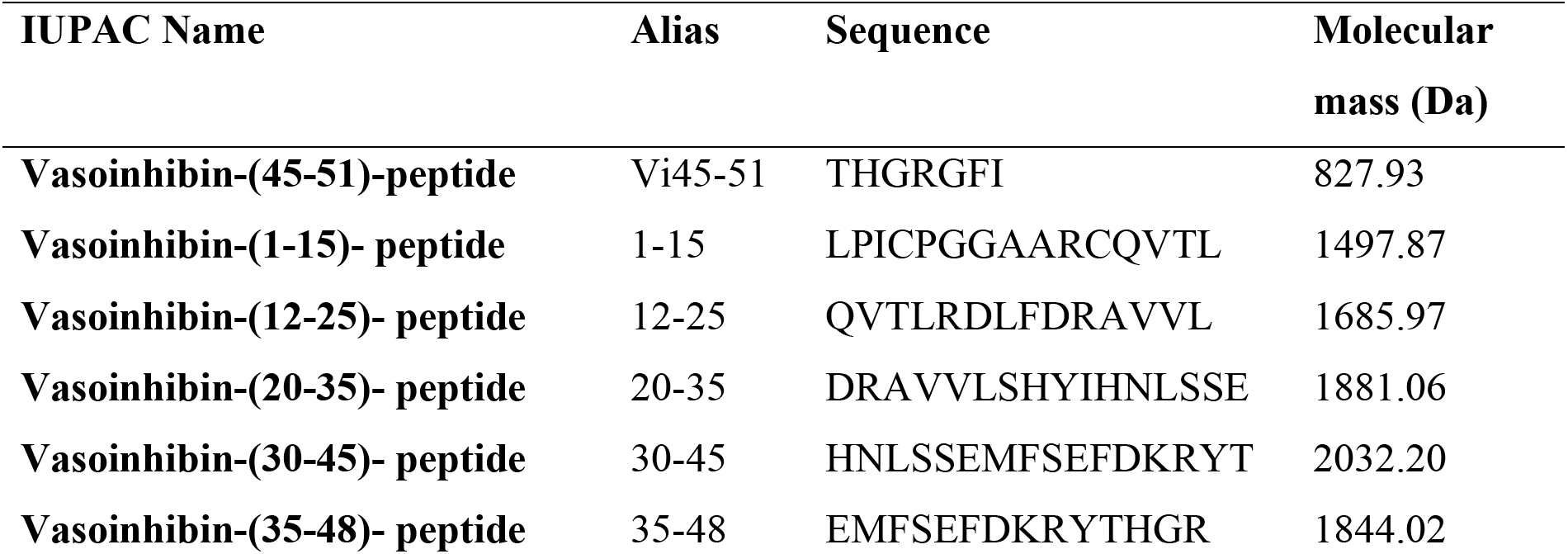
Synthetic oligopeptides.

### Cell culture

Human umbilical vein endothelial cells (HUVEC) were isolated (18) and cultured in F12K medium supplemented with 20% fetal bovine serum (FBS), 100 μg mL^-1^ heparin (Sigma Aldrich), 25 μg mL^-1^ endothelial cell growth supplement (ECGS) (Corning, Glendale, AZ), and 100 U mL^-1^ penicillin-streptomycin.

### Cell Proliferation

HUVEC were seeded at 14,000 cells cm^-2^ in a 96-well plate and, after 24 hours, starved with 0.5% FBS, F12K for 12 h. Treatments were added in 20% FBS, F12K containing 100 μg mL^-1^ heparin for 24 hours and consisted of 25 ng mL^-1^ VEGF and 20 ng mL^-1^ bFGF alone or in combination with 100 nM prolactin (PRL) (as negative control), 123-residue vasoinhibin (Vi1-123) or 48-residue vasoinhibin (Vi1-48) (positive controls), linear vasoinhibin analog (Vi45-51), cyclic retro-inverse-vasoinhibin analog (CRIVi45-51), synthetic oligopeptides mapping region 1 to 48 of vasoinhibin (1-15, 12-25, 20-35, 30-45, or 35-48). DNA synthesis was quantified by the DNA incorporation of the thymidine analog 5-ethynyl-2′-deoxyuridine (EdU; Sigma Aldrich) (10 μM) added at the time of treatments and labeled by the click reaction with Azide Fluor 545 (Sigma Aldrich) as reported (14,19). Total HUVEC were counterstained with Hoechst 33342 (Sigma Aldrich). Images were obtained in a fluorescence-inverted microscope (Olympus IX51, Japan) and quantified using CellProfiler software (20).

### Cell Invasion

HUVEC invasion was evaluated using the transwell matrigel barrier assay (21). HUVEC were seeded at 28,000 cells cm^-2^ on the luminal side of an 8-µm-pore insert of a 6.5 mm transwell (Corning) precoated with 0.38 mg mL^-1^ matrigel (BD Biosciences, San Jose, CA) in starvation medium (0.5% FBS F12K, without heparin or ECGS). Treatments were added inside the transwell and consisted of 100 nM PRL, Vi1-123, Vi1-48, Vi45-51, CRIVi45-51, or the oligopeptides 1-15, 12-25, 20-35, 30-45, or 35-48. Conditioned medium of 3T3L1 cells (ATCC, Manassas, VA) cultured for 2 days in 10% FBS was filtered (0.22 µm), supplemented with 50 ng mL^-1^ VEGF, and placed in the lower chamber as chemoattractant. Sixteen hours later, cells invading the bottom of the transwell were fixed, permeabilized, Hoechst-stained, and counted using the CellProfiler software (20).

### Leukocyte adhesion assay

HUVEC were seeded on a 96-well plate and grown to confluency. HUVEC monolayers were treated for 16 hours with 100 nM PRL, Vi1-123, Vi1-48, Vi45-51, CRIVi45-51, or the oligopeptides 1-15, 12-25, 20-35, 30-45, 35-48 in 20% FBS, F12K without heparin or ECGS. Treatments were added alone or in combination with anti-PAI-1 (5 μg mL^-1^), anti-uPAR (5 μg mL^-1^), or anti-β-tubulin (5 μg mL^-1^) antibodies. NF-κB activation inhibitor BAY 11-7085 (5 μM) was added 30 minutes prior to treatments. After the 16-hour treatment, HUVEC were exposed to a leukocyte preparation obtained as follows. Briefly, whole blood was collected into EDTA tubes, centrifuged (300 x *g* for 5 minutes), and the plasma layer discarded. The remaining cell pack was diluted 1:10 in red blood lysis buffer (150 mM NH_4_Cl, 10 mM NaHCO_3_, and 1.3 mM EDTA disodium) and rotated for 10 minutes at RT. The tube was centrifuged (300 x *g* 5 minutes), and when erythrocytes were no longer visible, leukocytes were collected by discarding the supernatant. Leukocytes were washed with cold PBS followed by another centrifugation step (300 x *g* 5 minutes) and resuspended in 5 mL of 5 μg mL^-1^ of Hoechst 33342 (Thermo Fisher Scientific) diluted in warm PBS. Leukocytes were incubated under 5% CO_2_-air at 37°C for 30 minutes, washed with PBS three times, and resuspended into 20% FBS, F12K to 10^6^ leukocytes mL^-1^. The medium of HUVEC was replaced with 100 μL of Hoechst-stained leukocytes (10^5^ leukocytes per well) and incubated for 1 hour at 37 °C. Finally, HUVEC were washed three times with warm PBS, and images were obtained in an inverted fluorescent microscope (Olympus IX51) and quantified using the CellProfiler software (20).

### Apoptosis

HUVEC grown to 80% confluency on 12-well plates were incubated under starving conditions (0.5 % FBS F12K) for 4 hours. Then, HUVEC were treated for 24 hours with 100 nM PRL, Vi1-123, Vi1-48, Vi45-51, CRIVi45-51, or the oligopeptides 1-15, 12-25, 20-35, 30-45, or 35-48 in 20% FBS, F12K without heparin or ECGS. Treatments were added alone or in combination with anti-PAI-1 (5 μg mL^-1^), anti-uPAR (5 μg mL^-1^), or anti-ϕ3-tubulin (5 μg mL^-1^) antibodies. NF-κB activation inhibitor BAY 11-7085 (5 μM) was added 30 minutes before treatments. Apoptosis was evaluated using the cell death detection ELISA kit (Roche, Basel, Switzerland). HUVEC were trypsinized, centrifuged, and resuspended with incubation buffer to 10^5^ cells mL^-1^. Cells were incubated at RT for 30 minutes and centrifugated at 20,000 x *g* for 10 minutes (Avanti J-30I Centrifuge, Beckman Coulter, Brea, CA). The supernatant was collected and diluted 1:5 with incubation buffer (final concentration ∼20^4^ cells mL^-1^). HUVEC concentration was standardized, and the assay was carried out according to the manufacturer’s instructions, measuring absorbance at 415 nm.

### Fibrinolysis assay

Human blood was collected into a 3.2% sodium citrate tube (BD Vacutainer) and centrifugated (1,200 x *g* for 10 minutes at 4 °C) to obtain plasma. Plasma (24 μL) was added to a 96-well microplate containing 20 μL of 50 mM CaCl_2_. Turbidity was measured as an index of clot formation by monitoring absorbance at 405 nm every 5 minutes after plasma addition. Before adding plasma, 0.5 μM of PAI-1 was preincubated in 10 mM Tris-0.01% Tween 20 (pH 7.5) at 37 °C for 10 minutes alone or in combination with 3 μM Vi1-123, Vi1-48, Vi45-51, CRIVi45-51, or the oligopeptides 1-15, 12-25, 20-35, 30-45, or 35-48. Once the clot was formed (∼20 minutes and maximum absorbance), treatments were added to a final concentration per well of 24% v/v plasma, 10 mM CaCl_2_, 60 pM human tissue plasminogen activator (tPA), 0.05 μM PAI-1, and 0.3 μM Vi1-123, Vi1-48, Vi45-51, CRIVi45-51, or the oligopeptides 1-15, 12-25, 20-35, 30-45, or 35-48. Absorbance (405 nm) was measured every 5 minutes to monitor clot lysis.

### PAI-1 binding assay

A 96-well ELISA microplate was coated overnight at 4 °C with 50 µL of 6.25 μM PRL, Vi1-123, Vi1-48, Vi45-51, CRIVi45-51, or the oligopeptides 1-15, 12-25, 20-35, 30-45, or 35-48, diluted in PBS. Microplate was blocked for 1 hour at RT with 5% w/v nonfat dry milk in 0.1% Tween-20-PBS (PBST), followed by three washes with PBST. Next, 100 nM of PAI-1 diluted in 0.2 mg mL^-1^ BSA-PBST was added and incubated for 1 hour at RT, followed by a three-wash step with PBST. Anti-PAI-1 antibodies (1 μg mL^-1^ diluted in blocking buffer) were added and incubated for 1 hour at RT. Microplates were then washed three times with PBST, and goat anti-rabbit HRP antibody (Jackson ImmunoResearch Labs, West Grove, PA, Cat# 111-035-144, RRID:AB_2307391) at 1:2,500 (diluted in 50% blocking buffer and 50% PBS) added and incubated for 1 hour at RT. Three last washes were done with PBST and microplates incubated for 30 minutes under darkness with 100 μL per well of an o-phenylenediamine dihydrochloride (OPD) substrate tablet diluted in 0.03% H_2_O_2_ citrate buffer (pH 5). Finally, the reaction was stopped with 50 µL of 3M HCl, and absorbance measured at 490 nm.

### NF-κB nuclear translocation assay

HUVEC were seeded on 1 μg cm^-1^ fibronectin-coated 18 mm-coverslips placed in 12-well plates and grown in complete media to 80% confluence. Then, cells were treated, under starving conditions (0.5 % FBS F12K), with 100 nM PRL, Vi1-123, Vi1-48, Vi45-51, CRIVi45-51, or the oligopeptides 20-35 or 30-45. After 30 minutes, cells were washed with PBS, fixed with 4% of paraformaldehyde (30 minutes), permeabilized with 0.5% Tx-100 in PBS (30 minutes), blocked with 5% normal goat serum, 1% BSA, 0.05% Tx-100 in PBS (1 hour), and incubated with 1:200 anti-NF-κB p65 antibodies (Santa Cruz Biotechnology, Santa Cruz, CA, Cat# sc-8008, RRID:AB_628017) in 1% BSA, 0.1% Tx-100 PBS overnight in a humidity chamber at 4 °C. HUVEC were washed and incubated with 1:500 goat anti-mouse secondary antibodies coupled to Alexa fluor 488 (Abcam, Cambridge, UK, Cat# ab150113, RRID:AB_2576208) in 1% BSA, 0.1% Tx-100 PBS (2 hours in darkness). Nuclei were counterstained with 5 μg mL^-1^ Hoechst 33342 (Sigma-Aldrich). Coverslips were mounted with Vectashield (Vector Laboratories, Burlingame, CA) and digitalized under fluorescence microscopy (Olympus IX51).

### Quantitative PCR of HUVECs

Eighty % confluent HUVEC in 6-well plates under starving conditions (0.5 % FBS F12K) were treated for 4 hours with 100 nM PRL, Vi1-123, Vi1-48, Vi45-51, CRIVi45-51, or the oligopeptides 1-15, 12-25, 20-35, 30-45, or 35-48. RNA was isolated using TRIzol (Invitrogen) and retrotranscribed with the high-capacity cDNA reverse transcription kit (Applied Biosystems). PCR products were obtained and quantified using Maxima SYBR Green qPCR Master Mix (Thermo Fisher Scientific) in a final reaction containing 20 ng of cDNA and 0.5 μM of each of the following primer pairs for human genes: *ICAM1* (5’-gtgaccgtgaatgtgctctc-3’ and 5’-cctgcagtgcccattatgac-3’), *VCAM1* (5’-gcactgggttgactttcagg-3’ and 5’-aacatctccgtaccatgcca-3’), *IL1A* (5’-actgcccaagatgaagacca-3’ and 5’-ttagtgccgtgagtttccca-3’), *IL1B* (5’-ggagaatgacctgagcacct-3’ and 5’ggaggtggagagctttcagt-3’), *IL6* (5’-cctgatccagttcctgcaga-3’ and 5’-ctacatttgccgaagagccc-3’), *TNF* (5’-accacttcgaaacctgggat-3’ and 5’-tcttctcaagtcctgcagca-3’) were quantified relative to *GAPDH* (5’-gaaggtcggagtcaacggatt-3’and 5’-tgacggtgccatggaatttg-3’). Amplification consisted of 40 cycles of 10 seconds at 95°C, 30 seconds at the annealing temperature of each primer pair, and 30 seconds at 72°C. The mRNA expression levels were calculated by the 2^-ΔΔCT^ method.

### In vivo vascular inflammation

Vascular inflammation was evaluated as previously reported (22). Briefly, female C57BL6 mice (8 weeks old) were injected intravenously (i.v.) with 16.6 μg of Vi45-51 or 40.7 μg of 30-45 in 50 μL of PBS to achieve ∼10 μM in serum. Controls were injected i.v. with 50 μL PBS. After 2 hours, animals were euthanized by cervical dislocation and perfused intracardially with PBS. A fragment of the lungs, liver, kidneys, and whole eyes were dissected and placed immediately in TRIzol reagent and retrotranscribed. The expression of *Icam1*, *Vcam1*, *Il1b*, *Il6*, and *Tnf* were quantified relative to *Gapdh* by qPCR as indicated for HUVEC using the following primer pairs for the mouse genes: *Icam1* (5’-gctgggattcacctcaagaa-3’ and 5’-tggggacaccttttagcatc-3’), *Vcam1* (5’-attgggagagacaaagcaga-3’ and 5’-gaaaaagaaggggagtcaca-3’), *Cd45* (5’-tatcgcggtgtaaaactcgtca-3’ and 5’-gctcaggccaagagactaacgt-3’), *Il1b* (5’-gttgattcaaggggacatta-3’ and 5’-agcttcaatgaaagacctca-3’), *Il6* (5’-gaggataccactcccaacagacc-3’ and 5’-aagtgcatcatcgttgttcataca-3’), and *Tnf* (5’-catcttctcaaaattcgagtgacaa-3’ and 5’-tgggagtagacaaggtacaaccc-3’).

### Joint inflammation

Male C57BL/6 mice (8 weeks old) were injected into the articular space of knee joints with vehicle (saline), or 87 pmol of Vi45-51 (72 ng) or 30-45 (176.8 ng) in a final volume of 10 μL saline. Twenty-four hours after injections, animals were euthanized in a CO_2_-saturated atmosphere. Joints were extracted, pulverized with nitrogen, RNA extracted, retrotranscribed, and the expression of mouse *Il1b*, *Il6*, and *Inos* (5’-cagctgggctgtacaaacctt-3’ and 5’-cattggaagtgaagcgtttcg-3’) quantified relative to *Gapdh* by qPCR as described above.

## Results

### Antiangiogenic HGR-containing vasoinhibin analogs are neither apoptotic, inflammatory, nor fibrinolytic

The linear-(Vi45-51) and cyclic retro-inverse- (CRIVi45-51) HGR-containing vasoinhibin analogs, like vasoinhibin standards of 123 residues (Vi1-123) and 48-residues (Vi1-48) (Figure 1a) inhibited the VEGF- and bFGF-induced proliferation of HUVEC (Figure 1b) and the VEGF-induced invasion of HUVEC (Figure 1c) without affecting the basal levels. These results are confirmatory of the antagonistic properties of HGR-containing vasoinhibin analogs (14) and served to validate their use to explore other vasoinhibin actions. PRL is not antiangiogenic (10) and was used as a negative control.

Contrary to the two vasoinhibin isoforms (Vi1-123 and Vi1-48), the HGR-containing vasoinhibin analogs failed to induce the apoptosis and inflammatory phenotype of HUVEC as well as the lysis of a fibrin clot (Figure 1d-g). Vi1-123 and Vi1-48, but not Vi45-51, CRIVi45-51, or PRL, stimulated the apoptosis of HUVEC revealed by DNA fragmentation measured by ELISA (Figure 1d), the adhesion of peripheral blood leukocytes to HUVEC monolayers (Figure 1e), and the *in vitro* lysis of a plasma clot (Figure 1f, g). Once the clot is formed (time 0), adding the thrombolytic agent tPA stimulates clot lysis, an action prevented by the coaddition of PAI-1. PAI-1 inhibition was reduced by Vi1-123 but not by Vi45-51, CRIVi45-51, or PRL (Figure 1f, g). Because binding to PAI-1 mediates the fibrinolytic properties of vasoinhibin (23), the binding capacity to PAI-1 was evaluated by adding PAI-1 to ELISA plates coated with or without PRL, Vi1-123, Vi45-51, or CRIVi45-51. The absorbance of the HRP-labeled antibody-PAI-1 complex increased only in the presence of Vi1-123 but not in uncoated wells and wells coated with the two HGR-containing vasoinhibin analogs or with PRL (Figure 1h).

These findings show that the HGR-containing vasoinhibin analogs lack the apoptotic, inflammatory, and fibrinolytic properties of vasoinhibin. The fact that PRL is not inflammatory, apoptotic, or fibrinolytic indicates that, like the antiangiogenic effect (14), these vasoinhibin properties emerge upon PRL cleavage.

### HGR-containing vasoinhibin analogs do not stimulate the nuclear translocation of NF-κB and the expression of inflammatory molecules in HUVEC

Because vasoinhibin signals through NF-κB to induce the apoptosis and inflammation of endothelial cells (11,12), we asked whether HGR-containing vasoinhibin analogs were able to promote the nuclear translocation of NF-κB and the expression of proinflammatory mediators in HUVEC (Figure 2). The distribution of NF-κB in HUVEC was studied using fluorescence immunocytochemistry and monoclonal antibodies against the p65 subunit of NF-κB (Figure 2a). Without treatment, p65 was homogeneously distributed throughout the cytoplasm of cells. Treatment with Vi1-123 or Vi1-48, but not with Vi45-51, CRIVi45-51, nor PRL resulted in the accumulation of p65 positive stain in the cell nucleus (Figure 2a) indicative of the NF-κB nuclear translocation/activation needed for transcription. Consistently, only vasoinhibin isoforms (Vi1-123 or Vi1-48) and not the HGR-containing vasoinhibin analogs nor PRL induced the mRNA expression of genes encoding leukocyte adhesion molecules [intercellular adhesion molecule 1 (*ICAM1*) and vascular cell adhesion molecule 1 *(VCAM1*)] and proinflammatory cytokines [interleukin -1α (*IL1A*), interleukin-1β (*IL1B*), interleukin 6 (*IL6*), and tumor necrosis factor α (*TNF*)] in HUVEC (Figure 2b). These findings show that HGR-containing vasoinhibin analogs are unable to activate NF-κB to promote gene transcription resulting in the apoptosis and inflammation of HUVEC. Furthermore, these results suggest that a structural determinant, different from the HGR motif, is responsible for these properties.

**Figure 2.**
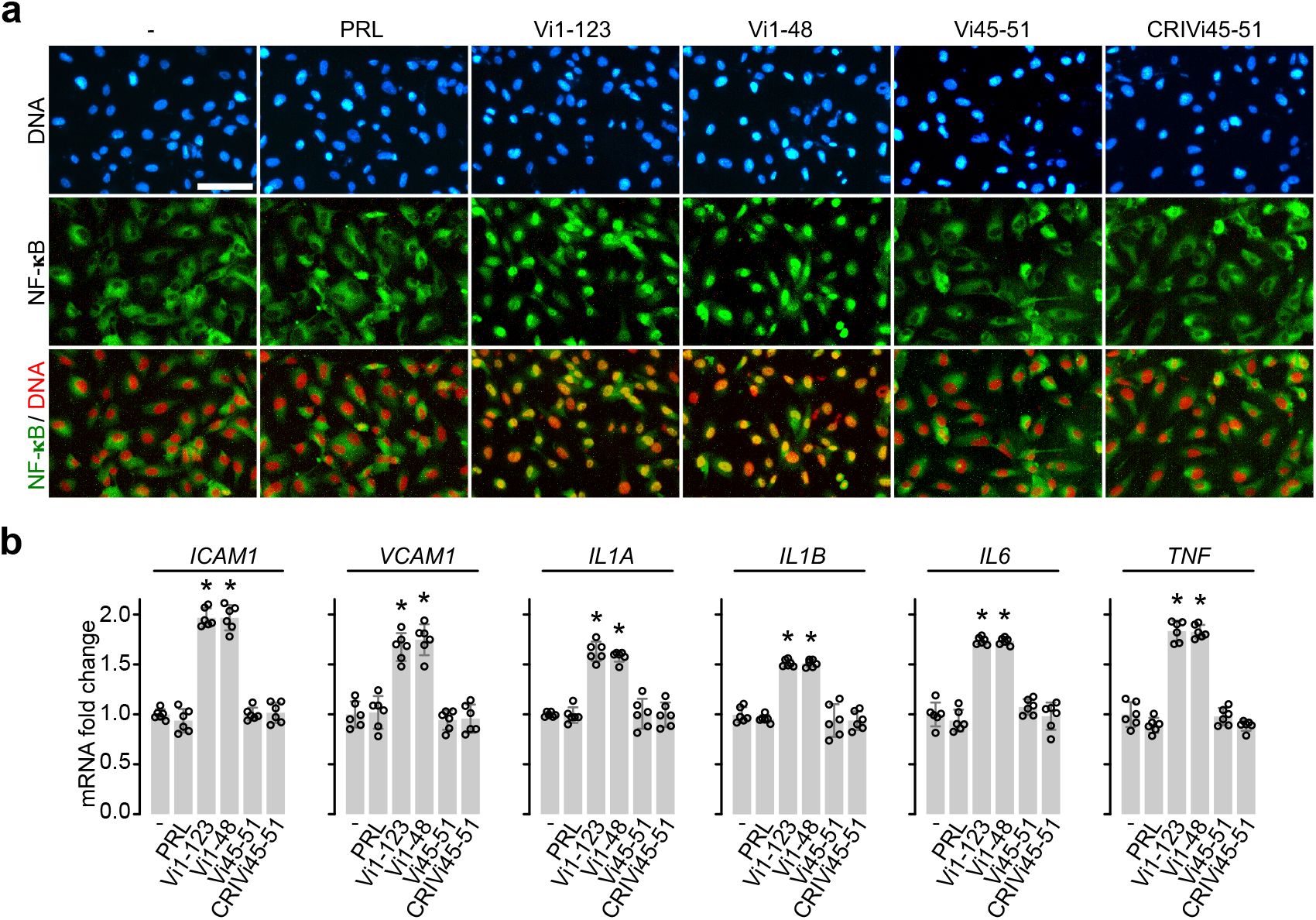
HGR-containing vasoinhibin analogs neither promote the nuclear translocation of NF-κB nor induce the expression of proinflammatory. **(a)** Immunofluorescence detection of NF-κB (p65) in HUVEC incubated in the absence (-) or presence of 100 nM of prolactin (PRL), the 123-residue (Vi1-123) and 48-residue (Vi1-48) vasoinhibin isoforms, and the linear (Vi45-51) and cyclic retro-inverse-(CRIVi45-51) HGR-containing vasoinhibin analogs. Scale bar = 100 µm. **(b)** HUVEC mRNA levels of cell adhesion molecules (*ICAM1* and *VCAM1*) and cytokines (*IL1A, IL1B, IL6,* and *TNF*) after incubation with or without (-) 100 nM of PRL, Vi1-123, Vi1-48, Vi45-51, or CRIVi45-51. Individual values are shown with open circles. Values are means ± SD of at least 3 independent experiments, *n* = 6, \**P* <0.001 vs. control without treatment (-) (One-way ANOVA, Dunnett’s).

### Oligopeptides containing the HNLSSEM vasoinhibin sequence are inflammatory, apoptotic, and fibrinolytic

Because the vasoinhibin of 48 residues (Vi1-48) conserves the apoptotic, inflammatory, and fibrinolytic properties of the larger vasoinhibin isoform (Vi1-123) (15), we scanned the sequence of the 48-residue isoform with synthetic oligopeptides (Figure 3a) for their ability to stimulate the apoptosis and inflammation of HUVEC and the lysis of a fibrin clot. First, we confirmed that only the oligopeptide containing the HGR motif (35–48) inhibited the proliferation and invasion of HUVEC, whereas the oligopeptides lacking the HGR motif were not antiangiogenic (Figure 3b, c).

**Figure 3.**
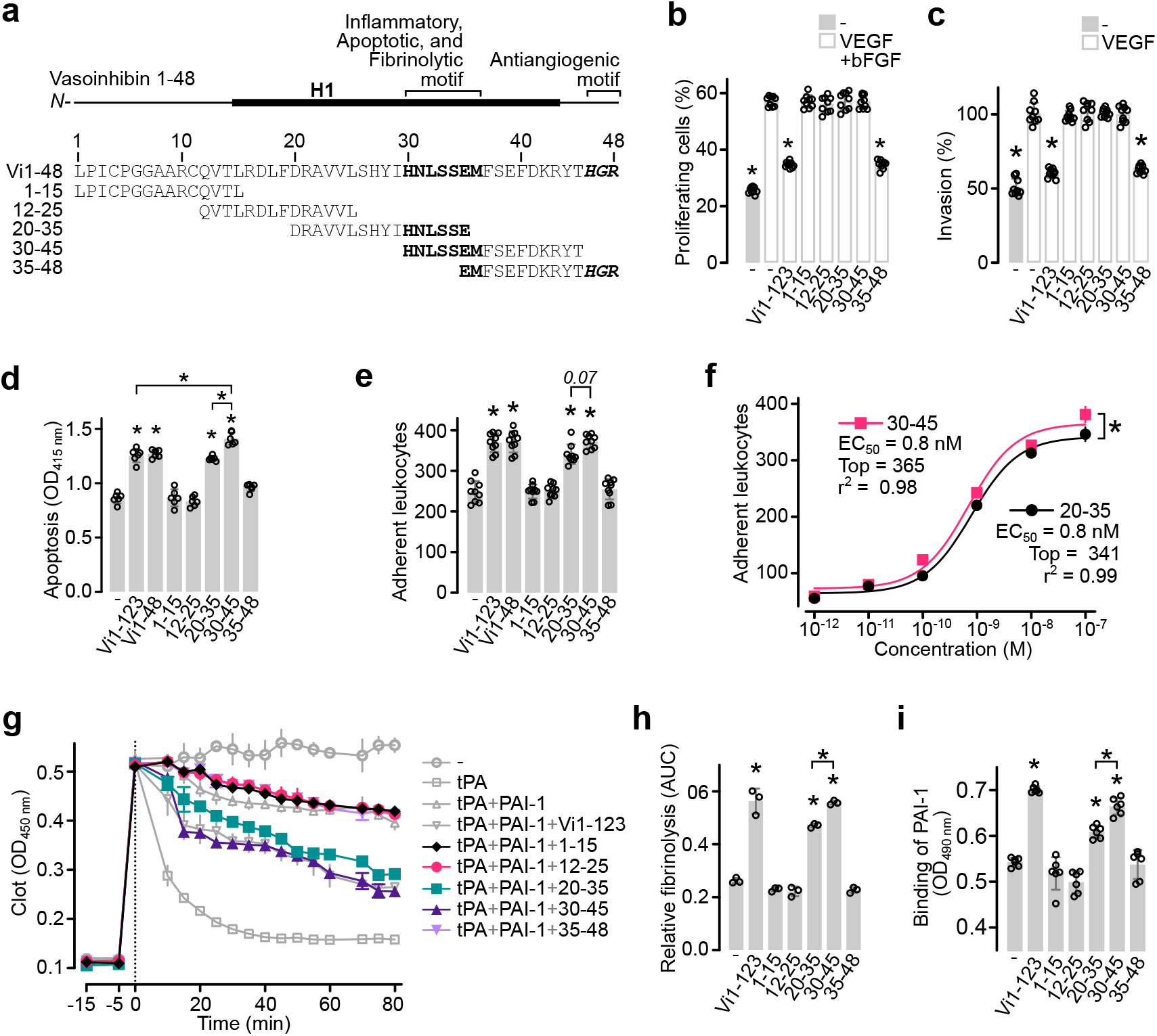
Oligopeptides containing the HNLSSEM sequence are inflammatory, apoptotic, and fibrinolytic. **(a)** Diagram of the sequence and localization of the oligopeptides scanning the 48-residue vasoinhibin isoform (Vi1-48). The α-helix 1 (H1), the inflammatory, apoptotic, and fibrinolytic sequence (HNLSSEM), and the antiangiogenic HGR motif are highlighted in bold. **(b)** Effect of 100 nM 123-residue vasoinhibin isoform (Vi1-123) or the scanning oligopeptides on the proliferation of HUVEC stimulated with 25 ng mL^−1^ of VEGF and 20 ng mL^−1^ of bFGF. Values are means ± SD relative to total cells (n=9). **(c)** Effect of 100 nM Vi1-123 or the scanning oligopeptides on the invasion of HUVEC stimulated with 25 ng mL^−1^ of VEGF. Values are means ± SD relative to VEGF stimulated values (*n = 9*). \**P*<0.0001 vs. VEGF+bFGF (-) or VEGF (-) controls. **(d)** Effect of 100 nM of Vi1-123, Vi1-48, or the scanning oligopeptides on HUVEC apoptosis (*n = 6*). **(e)** Effect of 100 nM of Vi1-123, Vi1-48, or the scanning oligopeptides on the leukocyte adhesion to a HUVEC monolayer (*n = 9*). **(f)** Dose-response of the leukocyte adhesion to HUVEC after treatment with the 20-35 or the 30-45 oligopeptides. Curves were fitted by least square regression analysis (*n = 6*, \**P* = 0.02, Paired T-test). **(g)** Lysis of a plasma clot by tissue plasminogen activator (tPA) alone or together with plasminogen activator inhibitor-1 (PAI-1) in the presence or absence of the scanning oligopeptides (*n = 3*). **(h)** Fibrinolysis relative to tPA and tPA+PAI-1 calculated with the area under the curve (AUC) of **(g)**. **(i)** Binding of PAI-1 to the immobilized scanning oligopeptides in an ELISA-based assay (*n = 6*). Individual values are shown with open circles. Values are means ± SD of at least 3 independent experiments, \**P* <0.001 vs. without treatment (-) controls (One-way ANOVA, Dunnett’s).

Only the oligopeptides 20-35 and 30-45 promoted the apoptosis of HUVEC (Figure 3d) and the leukocyte adhesion to HUVEC monolayers (Figure 3e, f) like Vi1-123 and Vi1-48. The estimated potency (EC_50_) of these oligopeptides was 800 pM, with a significantly higher effectiveness for the 30-45 (Figure 3f). Likewise, the 20-35 and 30-45 oligopeptides, but not oligopeptides 1-15, 12-35, or 35-48, exhibited fibrinolytic properties (Figure 3g, h) and bound PAI-1 like vasoinhibin (Figure 3i). The shared sequence between 20-35 and 30-45 oligopeptides corresponds to His30-Asn31-Leu32-Ser33-Ser34-Glu35 (HNLSSE) (Figure 3a). However, the significantly higher effect of 30-45 over 20-35 in apoptosis, inflammation, fibrinolysis, and PAI-1 binding, suggests that the Met36 could be a part of the apoptotic, inflammatory, and fibrinolytic linear determinant of vasoinhibin (HNLSSEM).

### Oligopeptides containing the HNLSSEM vasoinhibin sequence stimulate the nuclear translocation of NF-κB and the expression of inflammatory factors in HUVEC

Consistent with their apoptotic and inflammatory effects, the 20-35 and the 30-45 oligopeptides, like vasoinhibin (Vi1-123 and Vi1-48), induced the nuclear translocation of NF-κB (Figure 4a) and upregulated the mRNA expression levels of the leukocyte adhesion molecules (*ICAM1* and *VCAM1*) and inflammatory cytokines (*IL1A*, *IL1B*, *IL6*, and *TNF*) genes in HUVEC (Figure 5B).

**Figure 4.**
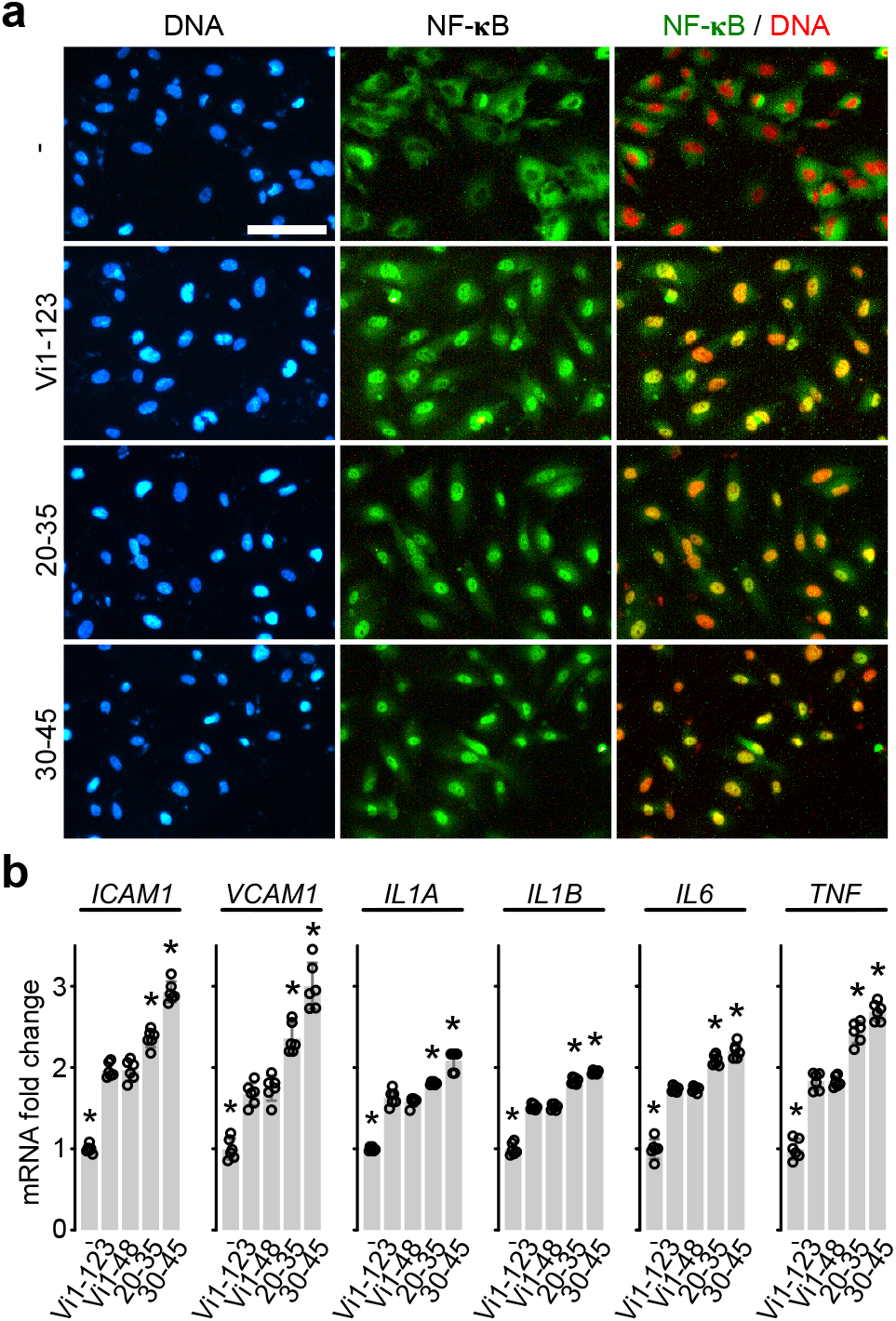
HNLSSEM-containing oligopeptides promote the nuclear translocation of NF-κB and induce the expression of proinflammatory genes. **(a)** Immunofluorescence detection of NF-κB (p65) in HUVEC incubated in the absence (-) or presence of 100 nM of the 123-residue vasoinhibin (Vi1-123), the 20-35 or the 30-45 residue-oligopeptides. Scale bar = 100 µm. Micrographs are representative of 3 independent experiments. **(b)** HUVEC mRNA levels of cell adhesion molecules (*ICAM1* and *VCAM1*) and cytokines (*IL1A, IL1B, IL6,* and *TNF*) after incubation with or without (-) 100 nM of Vi1-123, Vi1-48, 20-35, or 30-45. Individual values are shown with open circles. Values are means ± SD, *n* = 6, \**P* <0.001 vs. without treatment (-) controls (One-way ANOVA, Dunnett’s).

**Figure 5.**
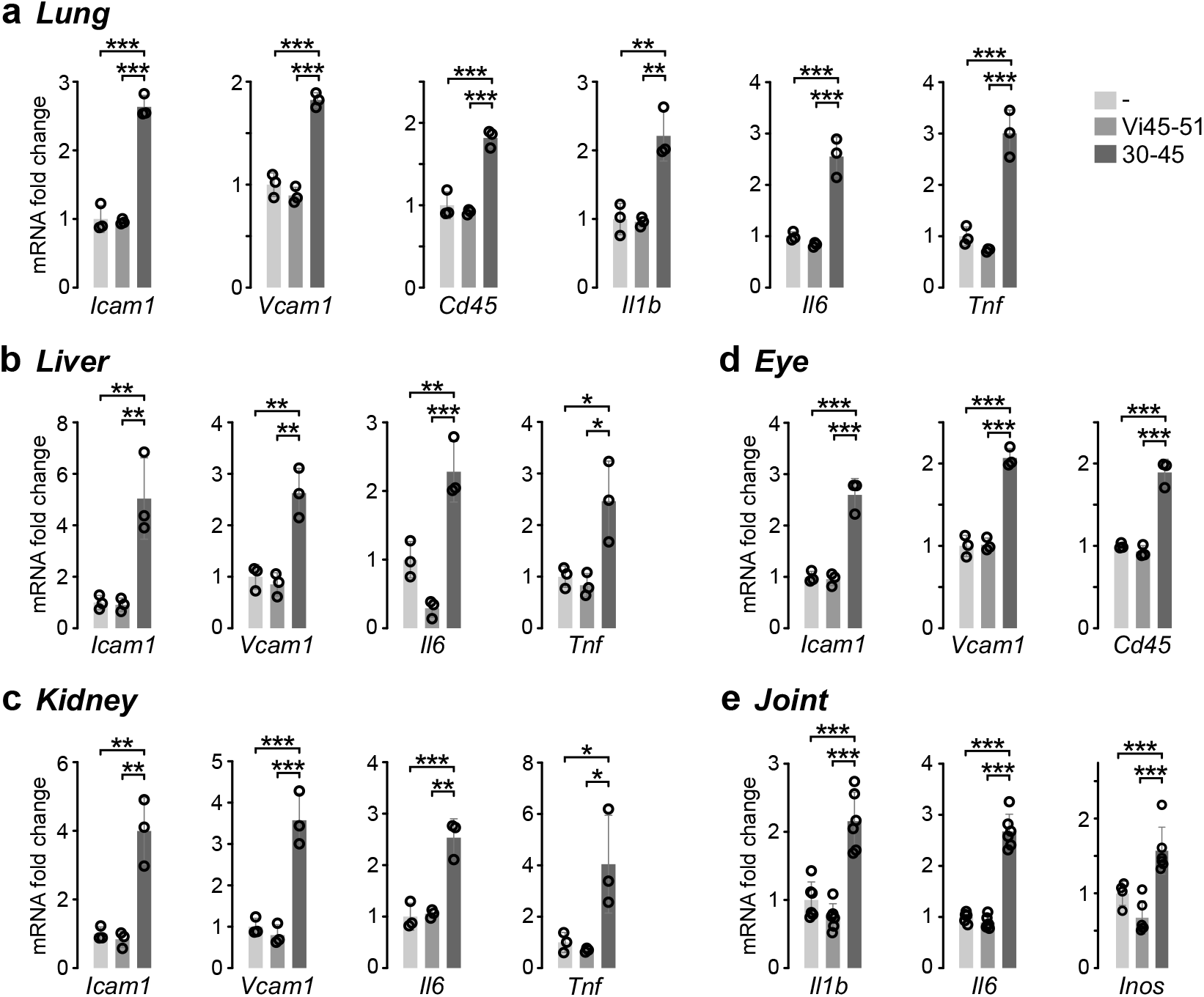
The HNLSSEM motif, but not the HGR motif, stimulates inflammation in vivo. Expression mRNA levels of adhesion molecules (*Icam1* and *Vcam1*), a leukocyte marker (*Cd45*), and cytokines (*Il1b*, *Il6*, and *Tnf*) in lung **(a)**, liver **(b)**, kidney **(c)**, and eye **(d)** from mice after 2 hours of intravenous injection of 16.6 μg of the HGR-containing vasoinhibin analog Vi45-51 or 40.7 μg of the HNLSSEM-containing 30-45 oligopeptide. **(e)** Expression mRNA levels of IL-1β, IL-6, and iNOS in knee joints of mice 24 h after the intra-articular injection of 72 ng of Vi45-51 or 176.8 ng of the 30-45 oligopeptide. Values are means ± SD, *n = 3*, \**P* <0.033, \*\**P* <0.002, \*\*\**P* <0.001 (One-way ANOVA, Tukey’s).

### In vivo inflammation is stimulated by the HNLSSEM sequence and not by the HGR motif

To evaluate whether the HGR or the HNLSSEM motifs promotes the inflammatory phenotype of endothelial cells *in vivo*, the HGR-containing vasoinhibin analog Vi45-51 or the HNLSSEM-containing 30-45 oligopeptide were injected i.v. to reach an estimated ≃10 μM concentration in serum, and after 2 hours, mice were perfused, and lung, liver, kidney, and eyes were collected to evaluate mRNA expression of leukocyte adhesion molecules (*Icam1* and *Vcam1*) and cytokines (*Il1b*, *Il6*, and *Tnf*), and the level of leukocyte marker (*Cd45*). The underlying rationale is that i.v. delivery and short-term (2-hour) analysis in thoroughly perfused animals would reflect a direct effect of the treatments on endothelial cell mRNA expression of inflammatory factors in the various tissues. The 30-45 peptide, but not the Vi45-51, increased the expression levels of these inflammatory markers in the evaluated tissues (Figure 5a-d). Furthermore, because vasoinhibin is inflammatory in joint tissues (24), we injected into the knee cavity of mice 87 pmol of the Vi45-51 or the 30-45 peptide, and after 24 hours, only the 30-45 oligopeptide induced the mRNA expression of *Il1b*, *Il6*, and inducible nitric oxide synthetase (*Inos*) (Figure 5e). The finding in joints implied that, like vasoinhibin, the inflammatory effect of the 30-45 peptide extends to other vasoinhibin target cells, i.e., synovial fibroblasts (24).

### PAI-1, uPAR, and NF-κB mediate the apoptotic and inflammatory effects of the HNLSSEM vasoinhibin determinant

Vasoinhibin binds to a multimeric complex in endothelial cell membranes formed by PAI-1, urokinase plasminogen activator (uPA), and uPA receptor (uPAR) (PAI-1-uPA-uPAR) (23), but it is unclear whether such binding influences vasoinhibin-induced activation of NF-κB, the main signaling pathway mediating its apoptotic and inflammatory actions (11,12,25). Because the HNLSSEM determinant in vasoinhibin binds to PAI-1, activates NF-κB signaling, and stimulates the apoptosis and inflammation of HUVEC, we investigated their functional interconnection by testing whether inhibitors of PAI-1, uPAR, or NF-κB modified the apoptosis and leukocyte adhesion to HUVEC treated with Vi1-123 and the oligopeptides 20-35 and 30-45 (Figure 6). Antibodies against uPAR and the inhibitor of NF-κB (BAY117085), but not the immunoneutralization of PAI-1, prevented the apoptotic effect of vasoinhibin and the HNLSSE-containing oligopeptides (Figure 6a). In contrast, all three inhibitors prevented the adhesion of leukocytes to HUVEC in response to Vi1-123, 20-35, and 30-45 (Figure 6b). These results indicate that vasoinhibin, through the HNLSSEM motif, uses PAI-1, uPAR, and/or NF-κB to mediate endothelial cell apoptosis and inflammation (Figure 6c).

**Figure 6.**
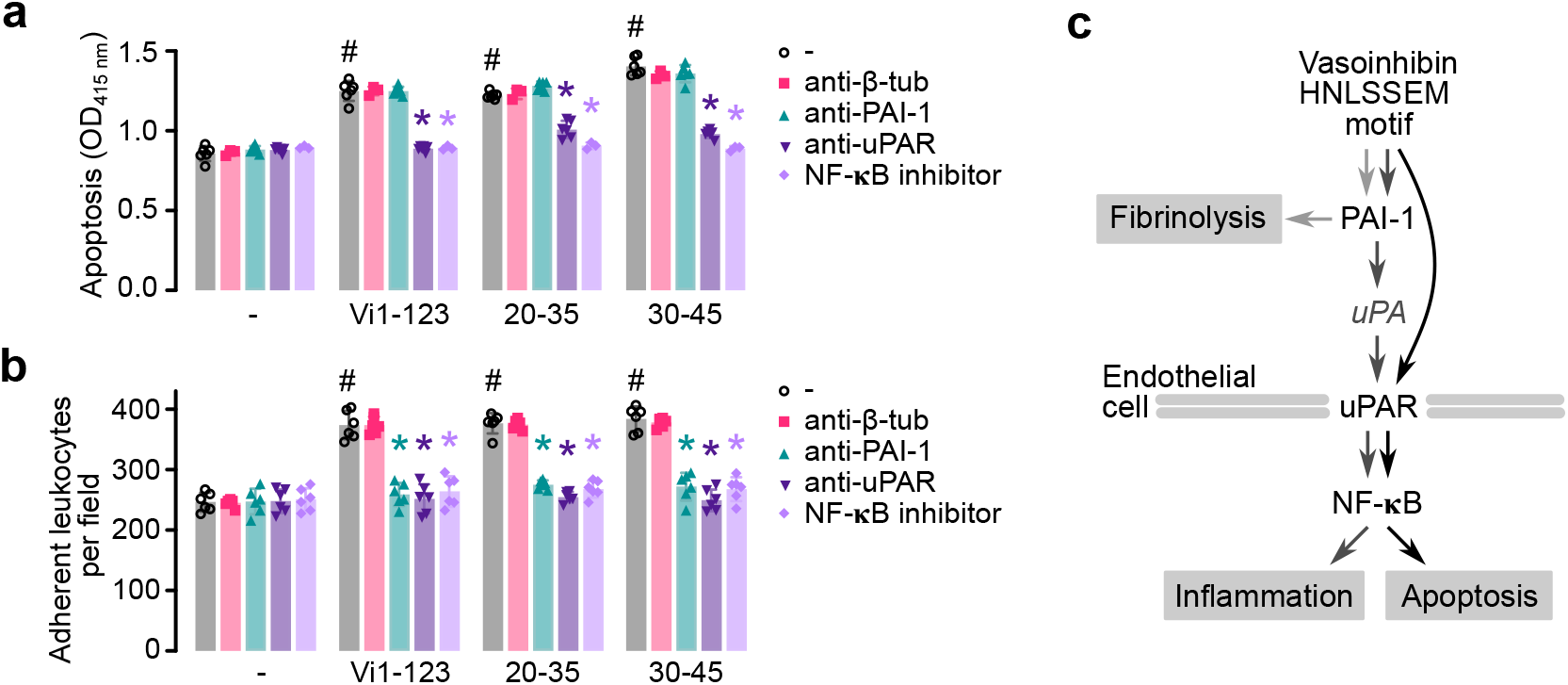
Inhibition of PAI-1, uPAR, and NF-κB signaling prevents the apoptotic and inflammatory effects of the HNLSSEM motif. Apoptosis of HUVEC **(a)** and leukocyte adhesion to a HUVEC monolayer **(b)** in response to the 123-residue vasoinhibin isoform (Vi1-123), the HNLSSEM-containing 20-35 or 30-45 oligopeptides in the absence (-) or presence of antibodies against β-tubulin (anti-β-tub), PAI-1 (anti-PAI-1) or uPAR (anti-uPAR), or the NF-κB inhibitor BAY117085. Values are means ± SD, *n ≥ 3*, # *P* <0.001 vs. respective untreated (-) group, \**P* <0.0001 vs. absence of antibodies (-) (Two-way ANOVA, Dunnett’s). **(c)** Schematic representation to illustrate that the HNLSSEM motif promotes fibrinolysis by binding to PAI-1 and endothelial cell apoptosis and inflammation via the activation of uPAR and NF-κB.

## Discussion

Vasoinhibin represents a family of proteins comprising the first 48 to 159 amino acids of PRL depending on the cleavage site of several proteases, including matrix metalloproteases (26), cathepsin D (27), bone morphogenetic protein 1 (28), thrombin (15), and plasmin (29). The cleavage of PRL occurs at the hypothalamus, the pituitary gland, and the target tissue levels defining the PRL/vasoinhibin axis (30). This axis contributes to the physiological restriction of blood vessels in ocular (31,32) and joint (26) tissues and is disrupted in angiogenesis-related diseases, including diabetic retinopathy (33), retinopathy of prematurity (34), peripartum cardiomyopathy (35), preeclampsia (36), and inflammatory arthritis (37). Furthermore, two clinical trials have addressed vasoinhibin levels as targets of therapeutic interventions (38).

However, the clinical translation of vasoinhibin is limited by difficulties in its production (39). These difficulties were recently overcome by the development of HGR-containing vasoinhibin analogs that are easy to produce, potent, stable, and even orally active to inhibit the growth and permeability of blood vessels in experimental vasoproliferative retinopathies and cancer (14). Nonetheless, the therapeutic value of HGR-analogs is challenged by evidence showing that vasoinhibin is also apoptotic, inflammatory, and fibrinolytic, properties that may worsen microvascular diseases (40,41). Here we show that the various functions of vasoinhibin are segregated into two distinct, non-adjacent, and independent small linear motifs: the HGR motif responsible for the vasoinhibin inhibition of angiogenesis and vasopermeability (14) and the HNLSSEM motif responsible for the apoptotic, inflammatory, and fibrinolytic properties of vasoinhibin (Figure 7).

**Figure 7.**
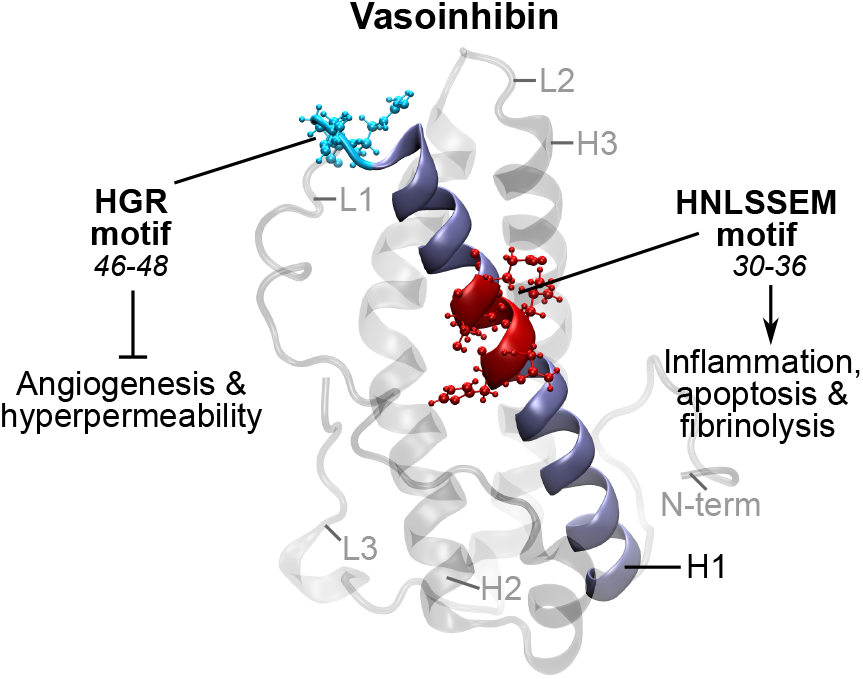
Vasoinhibin properties are segregated into two independent motifs. The vasoinhibin inhibitory properties on angiogenesis and vasopermeability are in the HGR motif (cyan), comprising residues 46 to 48, located in the early part of loop 1 (L1); whereas the inflammatory, apoptotic, and fibrinolytic properties of vasoinhibin reside in the HNLSSEM motif (red), comprising residues 30 to 36, located at the middle of α-helix 1 (H1) (navy blue). The vasoinhibin model was previously reported (43), and the figure was generated with Visual Molecular Dynamics software.

The HGR and HNLSSEM motifs are inactive in PRL, the vasoinhibin precursor. We confirmed that PRL has no antiangiogenic properties (10) and showed that PRL lacks apoptotic and inflammatory actions on endothelial cells as well as no fibrinolytic activity. PRL has 199 amino acids structured into a four-⍺-helix bundle topology connected by three loops (42). The HGR motif is in the first part of loop 1 (L1) connecting ⍺-helixes 1 and 2, whereas the HNLSSEM motif is in ⍺-helix 1 (H1) (Figure 7). Upon proteolytic cleavage, PRL loses its fourth ⍺-helix (H4), which drives a conformational change and the exposure of the HGR motif, obscured by H4 (14,43). Since H1 and H4 are in close contact in PRL (42), it is likely, that some elements of H4 also mask the HNLSSEM motif. Alternatively, it is also possible that residues of the HNLSSEM motif buried in the hydrophobic core of PRL become solvent exposed by the conformational change into vasoinhibin. However, this is unlikely since the hydrophobic core appears conserved during vasoinhibin generation (43).

A previous report indicated that binding to PAI-1 mediates the antiangiogenic actions of vasoinhibin (23). Contrary to this claim, antiangiogenic HGR-containing vasoinhibin analogs did not bind PAI-1, whereas the HNLSSEM-oligopeptides bound PAI-1 but did not inhibit HUVEC proliferation and invasion. While these findings unveil the structural determinants in vasoinhibin responsible for PAI-1 binding, they question the role of PAI-1 as a necessary element for the antiangiogenic effects of vasoinhibin. Little is known of the molecular mechanism by which vasoinhibin binding to the PAI-1-uPA-uPAR complex inhibits endothelial cells (23). Although the binding could help localize vasoinhibin on the surface of endothelial cells, the contribution of other vasoinhibin-binding proteins and/or interacting molecules cannot be excluded. For example, integrin ⍺5β1 interacts with the uPA-uPAR complex (44) and vasoinhibin binds to ⍺5β1 to promote endothelial cell apoptosis (45). Nevertheless, none of the HGR-containing analogs induced apoptosis. Therefore, the binding molecule/receptor that transduces the antiangiogenic properties of vasoinhibin remains unclear.

On the other hand, and consistent with previous reports (11,12,23), vasoinhibin binding to PAI-1 and activation of uPAR and NF-κB did associate with the apoptotic, inflammatory, and fibrinolytic properties of the HNLSSEM-containing oligopeptides. These oligopeptides, but not HGR-containing oligopeptides, induced endothelial cell apoptosis, nuclear translocation of NFκB, expression of leukocyte adhesion molecules and proinflammatory cytokines, and adhesion of leukocytes, as well as the lysis of plasma fibrin clot. The inflammatory action, but not the apoptotic effect, was prevented by PAI-1 immunoneutralization, whereas both inflammatory and apoptotic actions were blocked by anti-uPAR antibodies or by an inhibitor of NF-κB. Locating the apoptotic, inflammatory, and fibrinolytic activity in the same short linear motif of vasoinhibin is not unexpected since the three events can be functionally linked. The degradation of a blood clot is an important aspect of inflammatory responses, and major components of the fibrinolytic system are regulated by inflammatory mediators (46). Examples of such interactions are the thrombin-induced generation of vasoinhibin during plasma coagulation to promote fibrinolysis (15), the endotoxin-induced IL-1 production inhibited by PAI-1 (47), and the TNFα -induced suppression of fibrinolytic activity due to the activation of NFκB-mediated PAI-1 expression (48). Furthermore, uPA is upregulated by thrombin and inflammatory mediators in endothelial cells (49), and uPAR is elevated under inflammatory conditions (50).

The HNLSSEM-oligopeptides’ inflammatory action is further supported by their *in vivo* administration. The intravenous injection of HNLSSEM-oligopeptides upregulated the short-term (2-hour post-injection) expression of *Icam1* and *Vcam1*, *IlIb*, *Il6*, and *Tnf,* and the infiltration of leukocytes (evaluated by the expression levels of the leukocyte marker *Cd45*) in different tissues indicative of an inflammatory action on different vascular beds. Also, the HNLSSEM-oligopeptides injected into the intra-articular space of joints launched a longer-term inflammation (24-hour post-injection) indicative of an inflammatory response in joint tissues. This action is consistent with the vasoinhibin-induced stimulation of the inflammatory response of synovial fibroblasts, primary effectors of inflammation in arthritis (37).

The challenge is to understand when and how vasoinhibin impacts angiogenesis, apoptosis, inflammation, and fibrinolysis pathways under health and disease. One likely example is during the physiological repair of tissues after wounding and inflammation. By inhibiting angiogenesis, vasoinhibin could help counteract the proangiogenic action of growth factors and cytokines, whereas by stimulating apoptosis, inflammation, and fibrinolysis, vasoinhibin could promote the pruning of blood vessels, protective inflammatory reactions, and clot dissolution needed for tissue remodeling. However, in the absence of successful containment, overproduction of blood vessels, persistent inflammation, and dysfunctional coagulation determines the progression and therapeutic outcomes in cancer (41,51), diabetic retinopathy (52), and rheumatoid arthritis (53). The complexity of vasoinhibin actions under disease is exemplified in murine antigen-induced arthritis, where vasoinhibin ameliorates pannus formation and growth via an antiangiogenic mechanism but promotes joint inflammation by stimulating the inflammatory response of synovial fibroblasts (37,54).

Anti-angiogenic drugs, in particular VEGF inhibitors, have reached broad usage in the field of cancer and retinopathy, albeit with partial success and safety concerns (6,55,56). They display modest efficacy and survival times, resistance, and mild to severe side effects that include infections, bleeding, wound healing complications, and thrombotic events. Toxicities illustrate the association between the inhibition of blood vessel growth and multifactorial pathways influencing endothelial cell apoptosis, inflammation, and coagulation (6). The fact that the HGR-analogs lack the apoptotic, inflammatory, and fibrinolytic properties of vasoinhibin highlights their future as potent and safe inhibitors of blood vessel growth, avoiding drug resistance through their broad action against different proangiogenic substances.

In summary, this work segregates the activities of vasoinhibin into two linear determinants and provides clear evidence that the HNLSSEM motif is responsible for binding to PAI-1 and exerting apoptotic, inflammatory, and fibrinolytic actions via PAI-1, uPAR, and NF-κB pathways, while the HGR motif is responsible for the antiangiogenic effects of vasoinhibin. This knowledge provides tools for dissecting the differential effects and signaling mechanisms of vasoinhibin under health and disease and for improving its development into more specific, potent, and less toxic antiangiogenic, proinflammatory fibrinolytic drugs.

## Acknowledgments

We thank Xarubet Ruíz Herrera, Fernando López Barrera, Adriana González Gallardo, Alejandra Castilla León, José Martín García Servín, and María A. Carbajo Mata for their excellent technical assistance.

## Data Availability

Original data generated and analyzed during this study are included in this published article or in the data repositories listed in References.

